# The autophagy initiating kinase ULK1 is required for pancreatic cancer cell growth and survival

**DOI:** 10.1101/2021.05.15.444304

**Authors:** Sonja N. Brun, Gencer Sancar, Jan Lumibao, Allison S. Limpert, Huiyu Ren, Angela Ianniciello, Herve Tiriac, Michael Downes, Danielle D. Engle, Ronald M. Evans, Nicholas D.P. Cosford, Reuben J. Shaw

## Abstract

Amongst cancer subtypes, pancreatic ductal adenocarcinoma (PDA) has been demonstrated to be most sensitive to autophagy inhibition, which may be due to unique metabolic rewiring in these cells. The serine/threonine kinase ULK1 forms the catalytic center of a complex mediating the first biochemical step of autophagy. ULK1 directly recieves signals from mTORC1 and AMPK to trigger autophagy under stress and nutrient poor conditions. Studies in genetic engineered mouse models of cancer have revealed that deletion of core downstream autophagy genes (ATG5, ATG7) at the time of tumor iniation leads to a profound block in tumor progression leading to the development of autophagy inhibitors as cancer therapeutics. However, most preclinical studies and all clinical studies have relied on non-specific lysomotropic agents such as chloroquine and its derivatives, whose toxicity and off-target issues preclude further clinical development and which do not represent the impact of solely biochemically disrupting the autophagy pathway. Furthermore, druggable targets in the core autophagy pathway are quite limited, with ULK1 and ULK2 representing the only protein kinases in the pathway. Here we explore the genetic requirement for ULK1 and ULK2 in human PDA cancer cell lines and xenografts, and take advantage of new small molecule ULK1 inhibitors to demonstrate that ULK inhibition can overcome autophagy induction triggered by PDA therapeutics including chemotherapy and MEK inhibition. Finally we show that ULK inhibitors increase MHC Class I in PDA cells, suggestion a potential therapeutic avenue for such agents in combination with checkpoint immunotherapy.

## Introduction

Macroautophagy (henceforth referred to as autophagy) is a highly conserved pathway in all eukaryotic cells that mediates the coordinated lysosome-dependent degradation and recycling of cellular components (Levine and Kroemer, 2019). By degrading damaged organelles, misfolded proteins, and pathogens, autophagy maintains cellular homeostasis and promotes cell survival under stress (Amaravadi et al., 2019; Santana-Codina et al., 2017). Autophagic flux is maintained at a basal level in all cells, but it can be further increased by various stressors including deprivation of nutrients or exposure of cells to various chemical damaging agents. Biochemically, autophagy is a multistep pathway initiated through the ULK preinitiation complex, which consists of unc-51-like autophagy activating kinase 1 (ULK1) or its highly related homologue ULK2, in complex with three other proteins: autophagy related protein 13 (ATG13), ATG101 and focal adhesion kinase (FAK) family-interacting protein (FIP200) (Figure S1A). Autophagy is stimulated under low energy conditions by activation of AMP-activated protein kinase (AMPK) and inhibited under non-stressed conditions by the mechanistic target of rapamycin complex 1 (mTORC1) (Russell et al., 2014). mTORC1 inhibits autophagy by directly phosphorylating ULK1 on Ser757, as well as phosphorylating other components of the autophagy pathway. Conversely, AMPK activates ULK1 via two mechanisms: first, directly phosphorylating ULK1/2 at multiple sites N-terminal to the mTORC1 phosphorylation site, triggering ULK1/2 activation, and secondly by inhibiting mTORC1 activity via phosphorylation of the mTOR complex components Raptor and tuberous sclerosis complex 2 (TSC2) (Herzig and Shaw, 2018). Upon activation, ULK1/2 directly phosphorylate multiple components of the class III phosphatidylinositol 3-kinase [PI(3)KCIII] complex (Egan et al., 2015; Russell et al., 2013), consisting of VPS34, p150 VPS15, ATG14, and Beclin1 to nucleate autophagosome formation. Subsequently, the ATG9 transmembrane protein mediates the trafficking of source membrane— including from the endoplasmic reticulum, Golgi complex, mitochondria, endosome, and plasma membrane—for autophagosome elongation (Amaravadi et al., 2019). ULK1 has been reported to directly phosphorylate ATG9 (Mack et al., 2012; Papinski et al., 2014; Weerasekara et al., 2014), though the functional requirement for that event in autophagosome elongation or autophagosome vesicular trafficking in mammalian cells remains unclear. Next, two ubiquitin-like conjugation systems participate in autophagosome closure and maturation. A series of protein-to-lipid conjugation cascades attach a protein, an LC3 family member, to an autophagosome membrane lipid, which both identifies the vesicle as an autophagosome and facilitates the receipt of cargo. Both the ATG7–ATG3 and the ATG5–ATG12–ATG6L1 complex are required to conjugate LC3 (ATG8) family members (including GABARAPs) to the lipid phosphatidylethanolamine (PE), which is enriched in autophagosome membranes (Amaravadi et al., 2019; Levine and Kroemer, 2019). Meanwhile, in order for LC3 to be conjugated to lipid by this cascade, the cysteine protease ATG4 is required to process pro-LC3 to its soluble form (LC3-I). Once LC3 is conjugated to lipid, it becomes inserted on the surface of the emerging autophagosome. The lipidated form of LC3 (LC3-II) migrates faster than LC3-I on gel electrophoresis, allowing the ratio of lipidated to free LC3 to serve as an approximation of the number of AVs forming at any given time (Levine and Kroemer, 2019). Data from germline knockout of autophagy components in mice reveals that while ULK1 and ULK2 mice are each viable, the ULK1/2 double knockout mice are lethal soon after birth with a specific defect extremely similar to that reported in ATG5 mice (Cheong et al., 2014), suggesting that at least for these developmental processes, ULK1/2 are as essential as core downstream autophagy components. However, recent work reveals that kinases other than ULK, namely tank binding kinase 1 (TBK1), may promote the assembly of ATG13–FIP200 protein complex via phosphorylation of Syntaxin17 (Kumar et al., 2019), and other stress kinases besides ULK1 can also phosphorylate Beclin and Vps34 (Kim and Guan, 2013; Wei et al., 2015), raising the question of whether ULK1/2 are as essential for autophagy in all conditions as originally hypothesized.

Autophagy is critical for multiple aspects of cancer biology, including protein and organelle turnover, providing metabolites for cell survival and therapeutic resistance. Pancreatic ductal adenocarcinoma (PDA) is the most common form of sporadic pancreatic cancer. PDA cancer cells survive the harsh metabolic demands of rapid proliferation in a nutrient-poor tumor microenvironment (TME) through autophagy. Compared to other tumor types, PDA exhibits high levels of basal autophagy, thus rendering PDA tumors more sensitive to its inhibition compared to other cancers (Yang et al., 2014; Yang et al., 2011). Highlighting the importance of this pathway in PDA, perturbation of core autophagy components in PDA genetic engineered mouse (GEM) models via deletion of Atg5 or expression of a dominant negative mutant of Atg4B, blocks tumor progression (Yang et al., 2014; Yang et al., 2011). However a direct genetic examination of ULK1, ULK2 or any components of the ULK complex has not been performed in pancreatic cancer models to date. Importantly, autophagy not only helps PDA cancer cells survive under basal conditions, but is also critical as a survival mechanism chemotherapy, immunotherapy, and targeted therapies, such as MEK inhibition. Moreover, while poorly understood, accumulating evidence suggests that autophagy within the stromal compartment is critical for supporting tumor growth and therapeutic resistance (Sousa et al., 2016). Indeed, autophagy has been shown to be critical in both tumor cells and stroma in PDA, and a recently study demonstrated that hyperactive autophagy in PDA is responsible for downregulation of MHC Class I, contributing to the insensitivity of PDA to checkpoint immunotherapies(Yamamoto et al., 2020). Importantly, treatment of PDA cells with chloroquine or genetic inhibition of many different components of the autophagy pathway were capable of elevating cell surface MHC Class I and inducing therapeutic response to checkpoint immunotherapies (Yamamoto et al., 2020). Consequently, autophagy has emerged as an attractive therapeutic target for PDA. In fact, the autophagy inhibitor hydroxychloroquine has been tested in the clinic as a sensitizer to chemotherapy in PDA (Karasic et al., 2019; Mukubou et al., 2010; Wolpin et al., 2014). However, toxicity issues, lack of specificity of chloroquine derivatives, or limited bioavailability of currently available autophagy selective inhibitors have led to little clinical success (Limpert et al., 2018). To overcome these obstacles, we have developed the selective and small molecule ULK1/2 (ULKi) kinase inhibitors (Egan et al., 2015; Ren et al., 2020) and here we explore their use in PDA cells.

## Results

### Genetic ablation of ULk1 impairs growth and survival of pancreatic tumors similar to other autophagy regulators

Recent studies have identified that proteins in the autophagy pathway play a critical role in pancreatic tumor growth and progression (Amaravadi et al., 2019) but it remains unclear whether inhibiting the components in the initial autophagy pre-initiation (ULK1) complex would be as effective as loss of downstream autophagy machinery proteins (e.g. ATG5, ATG7) (see Figure S1A). To address this question, we generated a panel of knockout lines in human pancreatic cancer cell lines using CRISPR-based gene editing and multiple guides to target ULK1, ULK2, and both in combination. Additionally, for comparison, we generated knockouts of known autophagy regulators such as ATG5 and ATG7 with established roles in tumor growth. Western blots revealed efficient knockout of autophagy regulators in MiaPaca2 and Panc1 cell lines at the protein level (Figure 1A and Figure S1E) as well as significant reduction in ULK2 mRNA levels using RT-qPCR (Figure S1B, Figure S1F). Analysis of ULK1 substrates within the autophagy pathway revealed significant inhibition of ULK1 signaling in the ULK1 single and ULK1/2 double knockout lines. ATG13, a member of the ULK1 pre-initiation complex, as well as Beclin1, a member of the VPS34 autophagy initiation complex, are both established direct downstream substrates of ULK1 in the autophagy pathway (Egan et al., 2015; Russell et al., 2013). The clear decrease in phosphorylation of ATG13 and Beclin1 in the ULK knockout lines indicates impairment of the autophagy pathway. While loss of autophagy regulators had minimal effect on growth of pancreatic lines grown in full media in 2D, where autophagy activity would be predicted to be minimal (Figure S1G), growing knockout cell lines under more stressful low density conditions revealed the importance of this pathway on pancreatic cell line growth (Figure S1BC-S1D). Consistent with previous findings, loss of ATG5 and ATG7 decreased colony formation by 40% compared to control lines. Importantly, loss of ULK1 or ULK2 individually inhibited cell growth to a similar degree as ablation of core autophagy machinery and loss of both ULK1 and ULK2 further impaired cell line colony formation.

**Figure 1.**
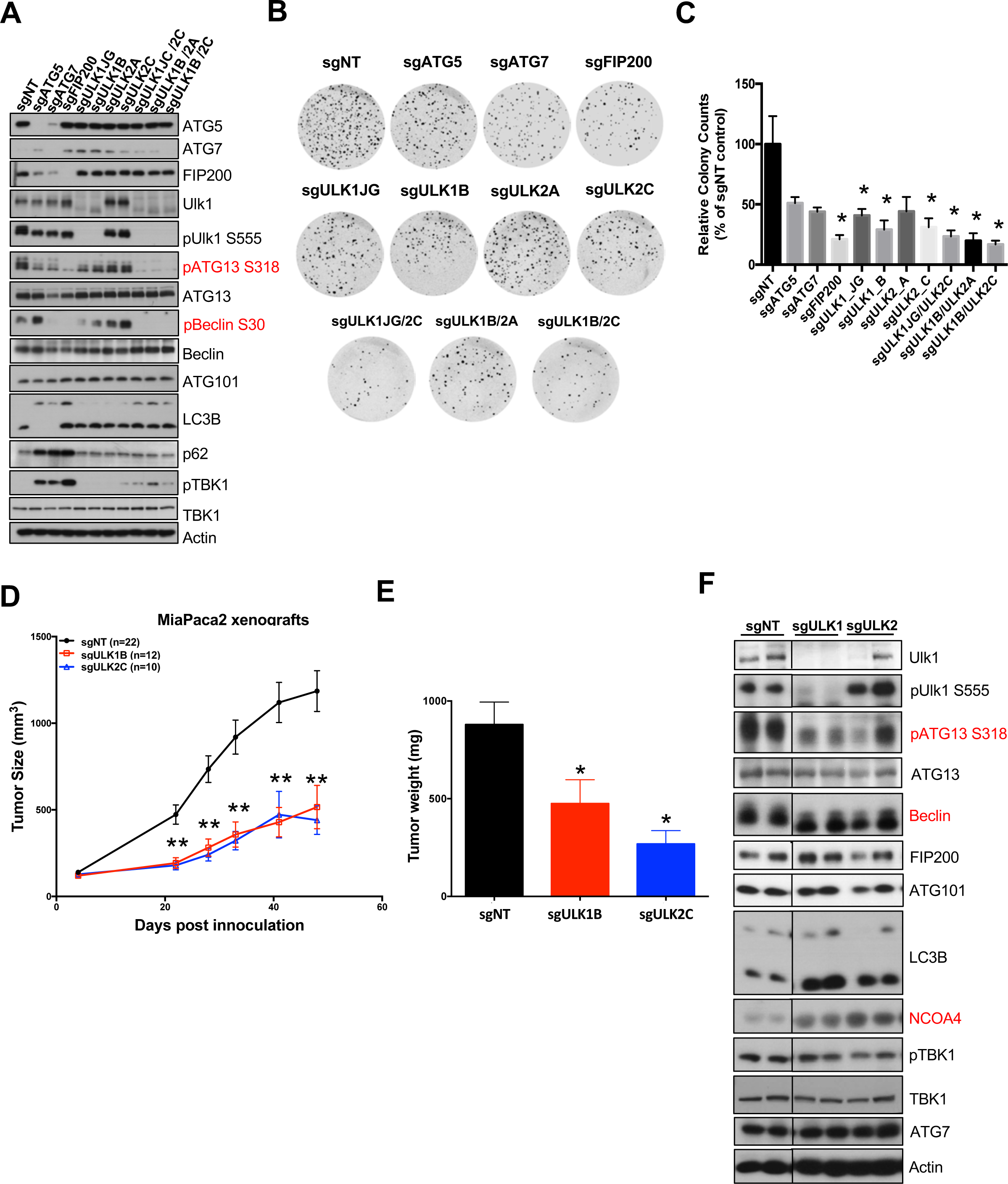
Loss of Ulk1 impairs growth and survival of PDAC cells similar to known autophagy regulators. **(A**) Western blots on lysates from MiaPaca2 cells stably expressing small guide RNAs (sg) targeting ULK1 or ULK2 alone, or lines targeted to co-delete ULK1 and ULK2 demonstrate efficient deletion. Direct ULK1 substrates are highlighted in red. Control sg-RNA treated MiaPaca2 cells (sgNT) as well as lines expressing guides targeting critical autophagy regulators ATG5, ATG7, and FIP200 are also shown for comparison. (***B-C***) Soft Agar analysis of MiaPaca2 CRISPR cell lines. (B) Representative images after 3 weeks of growth. Soft agar growth is most impaired in lines targeting both ULK1 and ULK2 concurrently. (C) Quantitation of average colony number expressed as a percentage of sgNT control cell colony number across three independent experiments plated in triplicate +/− SD. Significance relative to sgNT control was determined by one way Anova (p>0.0002) and t-test (*=p<0.05). **(D**) Caliper measurements of WT, sgULK1 or sgULK2 expressing MiaPaca2 xenografts plotted +/− SEM. Genetic CRISPR ablation of ULK1 or ULK2 decreases pancreatic tumor growth in MiaPaca2 xenograft models. Outliers were identified using ROUT outlier analysis with Q value of 1%, Significance determined relative to sgNT control tumor size by two-way Anova (p<0.001 time x genotype interaction) followed by one way Anova (p<0.0001) and t-test (**p<0.01) per timepoint. **(E)** Tumor weights of endpoint tumors from (D) +/− SEM. Significance determined relative to sgNT control tumor size by one way Anova (p<0.003) and t-test (p<0.04). (**F**) Tumors isolated from MiaPaca2 xenograft tumors were analyzed by western blotting for expression of autophagy proteins. Deletion of ULK1 and ULK2 results in decreased phosphorylation of ATG13 (direct ULK1 phosphorylation site), induces a mobility shift in Beclin, and increases NCOA4 expression.

To determine the ability of these cells to survive anoikis, which is associated with malignant invasion and metastasis, we then examined the ability of autophagy deficient cell lines to grow under anchorage independent conditions (Figure 1B-1C). Loss of ULK1 and ULK2 independently significantly inhibited the ability of pancreatic lines to form colonies in soft agar by over 50% compared to control lines using multiple guides (56-71% for ULK1 guides, 56-69% for ULK2 guides in MiaPaca2 cells). Colony numbers were also decreased through knockdown of ATG5 (49%) and ATG7 (56%), indicating that ULK1 knockdown impairs anchorage independent growth to a similar extent as loss of ATG proteins. Furthermore, ULK double knockout lines inhibited colony growth to a larger degree than any of the single autophagy knockouts (77-85%). Similar results were observed in Panc1 cells as well (Figure S1H-I). These data together suggest that loss of Ulk1 or ULK2 is as effective as loss of essential downstream regulators of autophagy on at preventing pancreatic cancer cell growth.

We next tested the importance of ULK proteins on *in vivo* tumor growth. MiaPaca2 ULK KO lines were injected into the flanks of nude mice and tumor growth monitored over time (Figure 1D). Loss of either ULK1 or ULK2 significantly impaired pancreatic flank xenograft growth and decreased tumor weights compared to control tumors (Figure 1E). Western blot analysis of the resulting tumor lysates demonstrated clear loss of ULK1 and decrease phosphorylation or bandshifts of direct ULK substrates in sgULK1 tumors. We also observed increases in LC3B II and NCOA4, indicating impairment of autophagy downstream of ULK1 loss (Figure 1F). While ULK2 KO tumors still express ULK1, they also displayed decreased phosphorylation of ATG13 and modulation of LC3B and NCOA4 similar to that observed in ULK1 KO tumors. Together, these data suggest that loss of ULK1 and ULK2 impairs autophagy and pancreatic tumor growth *in vivo*.

### ULK1 antagonists prevent autophagy, inhibit proliferation and promote apoptosis in pancreatic tumor cells

Having demonstrated the importance of ULK1 for growth of pancreatic tumor cells, we were next interested in evaluating the potential of ULK1 as a therapeutic target in cancer. To this end, we developed a novel potent and specific small molecule antagonist SBP-7455 to inhibit ULK1 activity (Ren et al., 2020). We first tested the ability of SBP-7455 to inhibit ULK1 in pancreatic cells in a dose-dependent manner by treating MiaPaca2 cells with drug for 1 hour, isolated lysates and examined phosphorylation ULK1 substrates by western blotting (Figure 2A). Treatment with SBP-7455 markedly decreased phosphorylation of ULK1 targets Beclin, ATG13, and ATG14. Notably, SBP-7455 inhibited ULK1 at a lower concentration than established ULK1 inhibitor SBI-0206965 and demonstrated less inhibition of off-targets such as FAK, indicating it is a more potent and specific inhibitor. Similar results were seen in both Panc1 and BxPC3 cells (Figure S2A), suggesting that SBP-7455 effectively inhibits Ulk1 activity in human pancreatic cancer cells.

**Figure 2.**
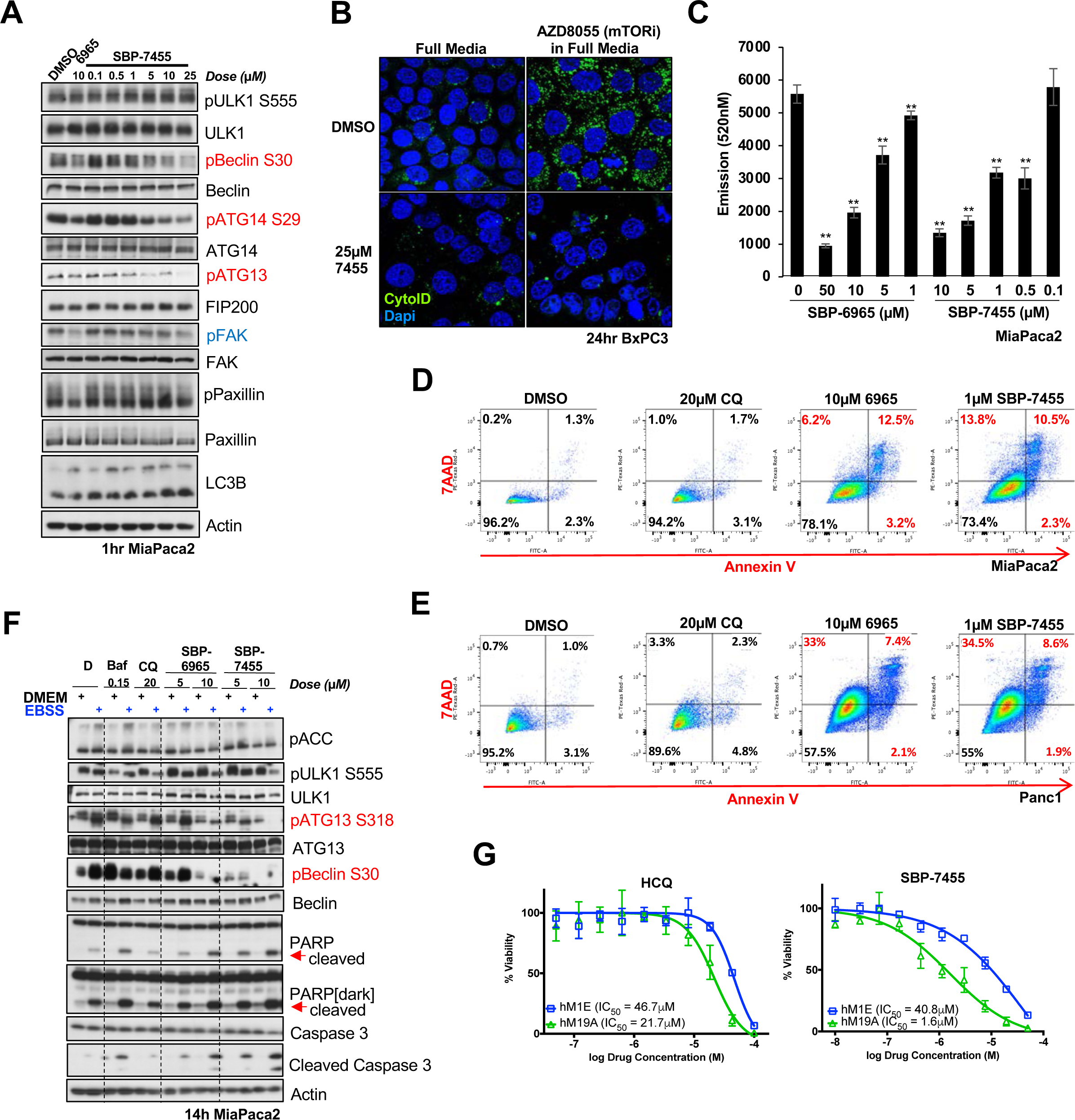
Pharmacological inhibition of ULK1/2 impairs survival of human PDAC cells. **A**) Lysates from MiaPaca2 cells treated with ULK1 inhibitors SBP-6965 and SBP-7455 at indicated concentrations for 1 hour were analyzed by immunoblotting for ULK1 inhibition and off-target signaling. SBP-7455 effectively inhibited phosphorylation of Ulk1 substrates Beclin, ATG13, and ATG14 with some inhibition of off-target FAK at high doses. **(B)** Representative immunofluorescent images of MiaPaca2 cells pre-treated with DMSO or 25μM SBP-7455 for two hours followed by 4μM AZD8055 (mTOR inhibitor) or combination of SBP-7455 and AZD8055 for 24hrs. Cells were stained with CytoID (green, autophagosome) and Dapi (blue, nuclei) to monitor autophagy vacuole formation. Pre-treatment with SBP-7455 prevents autophagosome formation stimulated by mTOR inhibition. **(C)** Dose dependent inhibition of MiaPaca cell growth by Ulk1 inhibitors SBP-6965 and SBP-7455 after 72hrs of treatment demonstrated using Cyquant assay to represent cell number run in triplicate in three independent experiments (representative run plotted +/− SD). SBP-7455 inhibits cell growth at lower concentrations than SBP-6965. Significance was determined by t-test (**p<0.01). **(D-E)** Miapaca2 (D) and Panc1 (E) cells treated for 72hrs with DMSO, 20μM chloroquine, 10μM SBP-6965, or 1μM 7455 were stained with AnnexinV and PI and analyzed by FACS. SBP-7455 induced apoptosis in pancreatic cells to a greater degree than autophagy inhibitor chloroquine or SBP-6965. **(F)** Lysates from MiaPaca2 cells treated with DMSO (D), Bafilomycin (Baf), chloroquine (CQ), SBP-6965, or SBP-7455 in full media or EBSS for 14hrs were analyzed for apoptosis induction by immunoblotting. Treatment with ULK1 inhibitors enhanced cleaved Parp and cleaved caspase signal induced by EBSS treatment to a greater degree than Baf or CQ treatment. **(G)** Two human organoid lines (hM1E and hM19A) were treated with a 10 concentrations of hydroxychloroquine or SBP-7455 for 5 days and dose response curves generated. Both lines responded in a dose-dependent manner to SBP-7455 with an IC50 of 40.8μM and 1.6μM respectively.

Given the established role of ULK1 in regulation of autophagy, we then tested the ability of SBP- 7455 to prevent autophagy initiation in pancreatic cancer cells. Pretreatment of cells with SBP-7455 significantly impaired formation of autophagosomes in response to stimulation by the mTOR inhibitor AZD8055 (Figure 2B). We then assessed the effects of autophagy inhibition through ULK1 on pancreatic tumor growth and survival. To address this, tumor cells were treated with our ULK1 inhibitors for 72 hours. We observed a dose dependent decrease in overall cell number after treatment with both SBI-0206965 or SBP-7455, with significant inhibition of cell growth at concentrations ≥0.5μM SBP-7455 (Figure 2C and Figure S2B-S2C). To determine if ULK1 inhibition also promotes cell death, tumor cells were treated with either chloroquine, a non-specific lysosomal inhibitor or our ULK1 inhibitors and then stained with Annexin-V and propidium iodide (PI). While inhibiting ULK1 significantly decreased the live cell percentage from 96.2% after DMSO treatment to 78.1 % (6965) and 73.4% (SBP-7455) in MiaPaca2 cells (Figure 2D), chloroquine (CQ) had almost no impact on apoptosis, suggesting that inhibition of ULK1 induces cell death more effectively. Similar results were seen in Panc1 cells (Figure 2E). Additionally, western blots of treated cells showed that autophagy inhibition using Bafilomycin (vacuolar ATPase inhibitor that prevents fusion of autophagosomes with lysosomes), SBI-0206965, and SBP-7455 enhances starvation-induced cleaved Parp and cleaved caspase 3 much more effectively than chloroquine (Figure 2F). Together, these data show that ULK1 inhibition is not merely cytostatic, but can also promote apoptosis of tumor cells.

Given the importance of ULK1 and autophagy in established pancreatic tumor cell lines, we were interested whether inhibition of autophagy and ULK1 would be effective in primary pancreatic cancer patient-derived organoids (PDOs). PDOs have been demonstrated to recapitulate the molecular diversity of pancreatic cancer patients and are being evaluated as a tool to predict patient drug response (Boj et al., 2015; Tiriac et al., 2018). IC50 values were calculated from two independent human organoid line dose response curves using hydroxychloroquine (HCQ), a chloroquine derivative currently in clinical trials, and SBP-7455 (Figure 2G and Figure S2D). hM1E is an organoid line established from a lung metastasis of a patient with KRAS^G12D^ mutant, classical pancreatic cancer while hM19A was established from a liver metastasis of a patient with KRAS^G12V^ mutant, basal-like pancreatic cancer (Tiriac et al., 2018). While the IC50 range for HCQ was 20-46 μM, cells were responsive to ULK1 inhibition by SBP-7455 with IC50 values up to ten-fold lower μM than those of HCQ in the hM19A organoids (21.7 μM HCQ vs 1.6 μM SBP-7455), indicating that human pancreatic cancer PDOs are sensitive to ULK1 antagonists, which are more effective than HCQ.

### Chemotherapy and targeted inhibitors induce autophagy which can be reversed using ULK1 inhibitors

Numerous studies have found that autophagy contributes to cancer cell resistance to first line therapies (Amaravadi et al., 2011; Levy et al., 2017; Qadir et al., 2008; Samaddar et al., 2008; Thorburn et al., 2014). Given this, we hypothesized that ULK1 inhibitors would reverse this resistant by preventing drug-induced autophagy upregulation and could synergize with established therapies to enhance response. To test this, we generated pancreatic cells expressing mCherry-GFP-LC3B to monitor autophagy flux in response to standard therapies and ULK1 inhibitors. As previously described (Castillo et al., 2017), LC3B incorporated into phagophores or autophagosomes express both mCherry and GFP equally, but after fusion with autolysosomes GFP is degraded in the low pH leaving only mCherry signal. Thus, autophagy flux can be monitored over time through examination of GFP/mCherry ratios. As expected, autophagy was robustly induced in MiaPaca2 and BxPC3 reporter lines were treated with mTOR inhibitor INK128 (Figure 3A-3B, S3A-C). Importantly, we observed a dose dependent rescue of autophagy induction when cells were treated with SBP-7455 indicating that ULK1 inhibitor treatment prevents mTOR induced autophagy. Furthermore, INK128 and SBP-7455 cooperate to significantly impair cell growth to a greater degree than either inhibitor alone, suggesting that SBP-7455 treatment enhances sensitivity to mTOR inhibitors (Figure S3D-E). We next asked whether autophagy induction is a mechanism of resistance to other standard chemo- and targeted therapies that can be reversed through use of SBP-7455. Treatment of MiaPaca2 reporter cells with Gemcitabine, Cisplatin, and the MEK inhibitor Trametinib all resulted in robust induction of autophagy (Figure 3C, S3F), which could be partially rescued by pre-treatment with SBP-7455 (Figure 3D, S3G). Together, these data suggest that treatment with ULK inhibitors can enhance response to multiple standard therapies.

**Figure 3.**
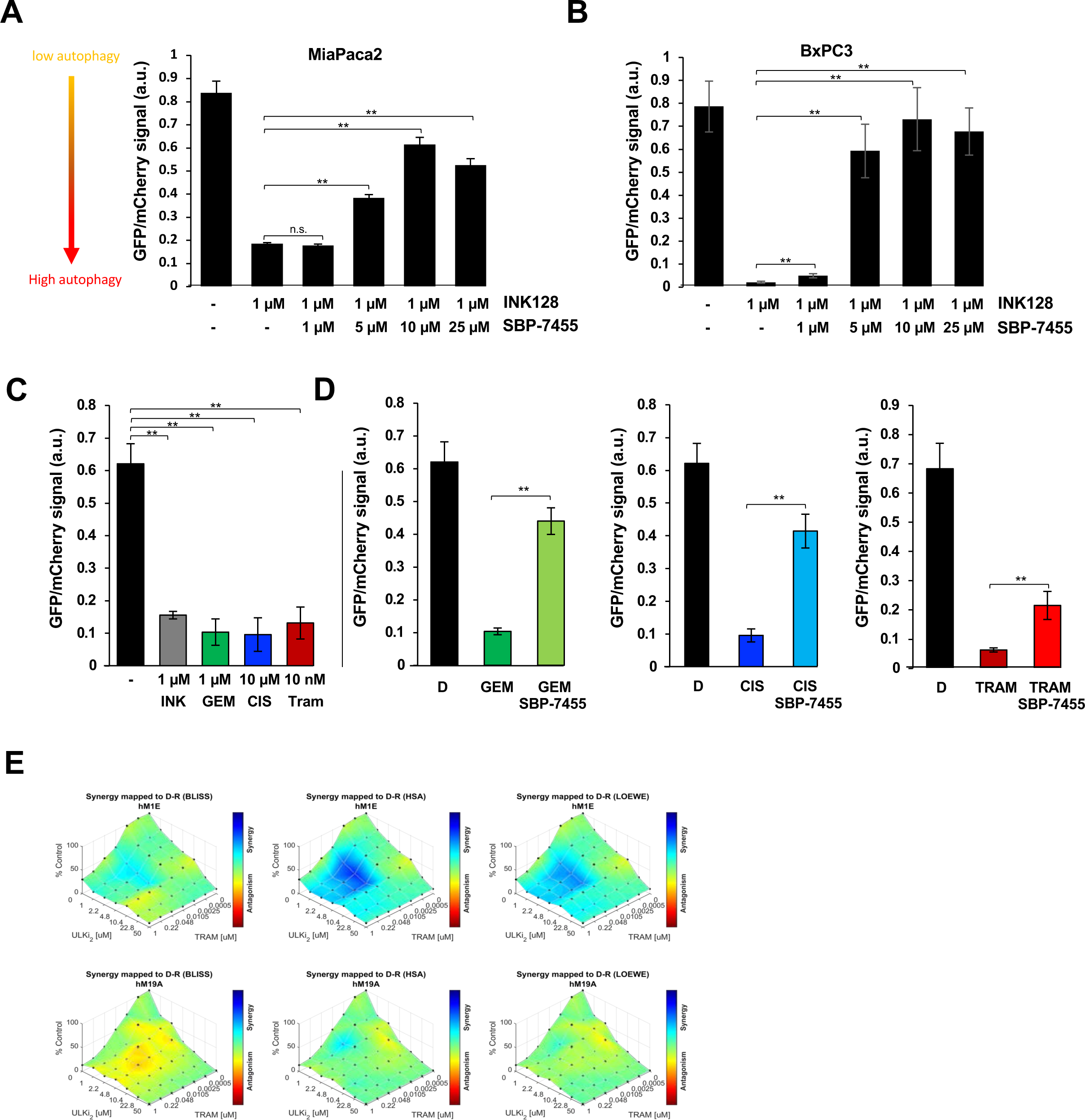
PDAC therapies induce autophagy, which is reversed by ULK1/2 inhibitors. **(A-B)** MiaPaca2 (A) and BxPC3 (B) cells stably expressing mCherry-GFP-LC3 construct treated with DMSO or SBP-7455 for 2 hours then 1 μM INK128 (mTOR inhibitor) was added to the cells in combination with ULK1 inhibitor as indicated and were monitored for GFP and mCherry signal using Incucyte fluorescent imaging over 36 hours to track autophagy induction. Plots of GFP/mCherry signal ratios at 36hrs (top) were generated by quantification of five wells per condition (22-36 pictures per condition +/− SEM). mTOR inhibitor robustly induces autophagy which is partially rescued by cotreatment with SBP-7455. Significance was determined by unpaired t-test (**=p>0.01). **(C-D)** MiaPaca2 cells stably expressing LC3-mCherry-GFP construct were treated wtih DMSO, 1 μM INK128 (INK) 1 μM Gemcitabine (GEM), 10 μM Cisplatin (CIS), and 10 nM Trametinib (Tram) either alone (C) or in combination with 10μM SBP-7455 (D) and GFP/mCherry ratio monitored over 36 hours using Incucyte fluorescent imaging to track autophagy flux. Plots of GFP/mCherry signal ratios at 33-36hrs were generated by quantification of 20x images of five wells per condition (34-35 pictures/condition +/− SEM). Chemotherapies and targeted therapies used to treat pancreatic cancer induce autophagy that is partially rescued by co-treatment with SBP-7455. Significance was determined by unpaired t-test (**=p>0.01). **(D)** Synergy matrices were generated using Combenefit for human organoids hM1E and hM19A treated with Trametinib and SBP-7455 for 5 days. SPB-7455 synergizes strongly with Trametinib in hM1E organoids and additive in hM19A organoids.

Having identified cooperation between autophagy inhibition using SBP-7455 and standard of care therapies in established pancreatic cell lines, we next explored the synergy between SBP-7455 and Trametinib in pancreatic cancer PDOs. Recent publications in *Nature Medicine* (Bryant et al., 2019; Kinsey et al., 2019) have highlighted the potential of autophagy inhibitors to be used to increase response to therapies targeting KRAS effector pathways such as blockade of MEK and/or ERK. These data, along with our own observations of autophagy induction in response to trametinib (Figure 3C and Figure S3F; rightmost panel), we decided to test efficacy of trametinib in combination with SBP-7455. Both human PDO lines exhibited similar responses to Trametinib (Figure S3H). While only additive effects were observed for hM19A, we observed clear synergy between Trametinib and SBP-7455 in hM1E organoids (Figure 3D). These studies show diversity of PDO response to SBP-7455 and Trametinib combination treatment, suggesting that comprehensive analyses of the range of responses and identification of predictive biomarkers are warranted in future studies. The strong synergy observed between SBP-7455 and trametinib indicates that for some patients, this combination represents a promising intervention approach.

### SBP-7455 regulates MHC1 expression in pancreatic tumor cells

In addition to combination of autophagy inhibition with standard therapies, recent studies highlighted the exciting potential for autophagy inhibitors to enhance response to immune therapies (Yamamoto et al., 2020). They observed that MHC class I molecules are predominantly localized in autophagosomes and lysosomes in pancreatic tumor cells, impairing antigen presentation, and that inhibition of autophagy using bafilomycin or cholorquine can cause accumulation and redistribution of these molecules. Given these implications, we asked whether more specific inhibition of autophagy through ULK1 can also alter MHC1 expression. Excitingly, western blotting revealed that accumulation of MHC-I in both Miapaca2 and BxPC3 cells in response to both 6965 and SBP-7455 (Figure 4A,B). Importantly, MHC-I expression was induced to a higher degree in SBP-7455 treated cells than those treated with CQ, suggesting that ULK inhibitors may be more effective than the currently available autophagy inhibitors for immune checkpoint blockade combination strategies. Using immunofluorescence techniques, we also examined colocalization of Lamp1 and MHC1 molecules in pancreatic cancer cells. Consistent with previous studies, we determined that MHC-I was primarily localized in Lamp1+ puncta under normal conditions (Figure 4C). Critically, in response to SBP-7455 treatment, we observed that the majority of MHC-1 signal was no longer overlapping with Lamp1. These data together suggest that SBP-7455 effectively increases MHC-I expression and promotes release from lysosomes, which has great potential to enhance antigen presentation and combine with immunotherapies to treat pancreatic cancer.

## Discussion

CRISPR-mediated disruption of ULK1 and ULK2 or their combination in MiaPaca2 and Panc1 human PDA cell lines demonstrates unequivocally that ULK1/2 depletion causes as profound a block to PDA cell growth in low density, soft-agar, and in flank xenografts, unlike the minimal effects observed in cells growing in full serum in 2D. These data fit with recent studies performed utilizing real-time imaging to track the fate of different tumor cell lines following CRISPR deletion of a panel of autophagy components (Towers et al., 2019; Towers et al., 2020), though pancreatic cancer cells were not examined in those studies. These data suggest that in conditional deletion of ***Ulk1*** and ***Ulk2*** alleles within pancreatic tumor cells may mimic the phenotype of Atg5 or Atg7 deletion in the KPC genetic engineered model of pancreatic adenocarcinoma, which will be a goal for future work. In addition to the unique sensitivity of pancreatic cancer cells to upregulation of autophagy for survival, recent studies indicate that the upregulated autophagy in PDA cells also results in the sequestration of MHC Class I in a cytoplasmic autophagosome/ lysosomes in these cells resulting in attenuated immune recognition of these tumor cells (Yamamoto et al., 2020). Remarkably, treatment of these cells with chloroquine or siRNA or sgRNA to a number of autophagy components, including ULK1, resulted in upregulation of MHC Class I protein levels on the cell surface (Yamamoto et al., 2020). We therefore examined here whether our ULK inhibitors were capable of regulating endogenous MHC Class I and find that indeed, a dose-dependent manner proportional to their inhibition of ULK1, these ULK kinase inhibitors led to increased MHC Class I (Figure 4). Taken together with recent studies that autophagy plays critical roles in the tumor microenvironment (Sousa et al., 2016), these data collectively support further exploration of ULK inhibitors as PDA therapeutics. A recent study reports the first data with a bioavailable ULK kinase inhibitor MRT-69882, which showed striking results in PDA xenografts, and synergized well with a tool compound that inhibited micropinocytosis, an alternative process for PDA cells to obtain critical metabolites when autophagy is blocked (Su et al., 2021). Our data with conventional therapeutics (gemcitabine) and targeted therapeutics (MEKi) (Figure 3), and the potential for immunotherapeutic combinations (Figure 4), makes a very unique modality which may be able to be combined with distinct current standards of care in the future treatment of PDA (Maitra, 2020).

**Figure 4.**
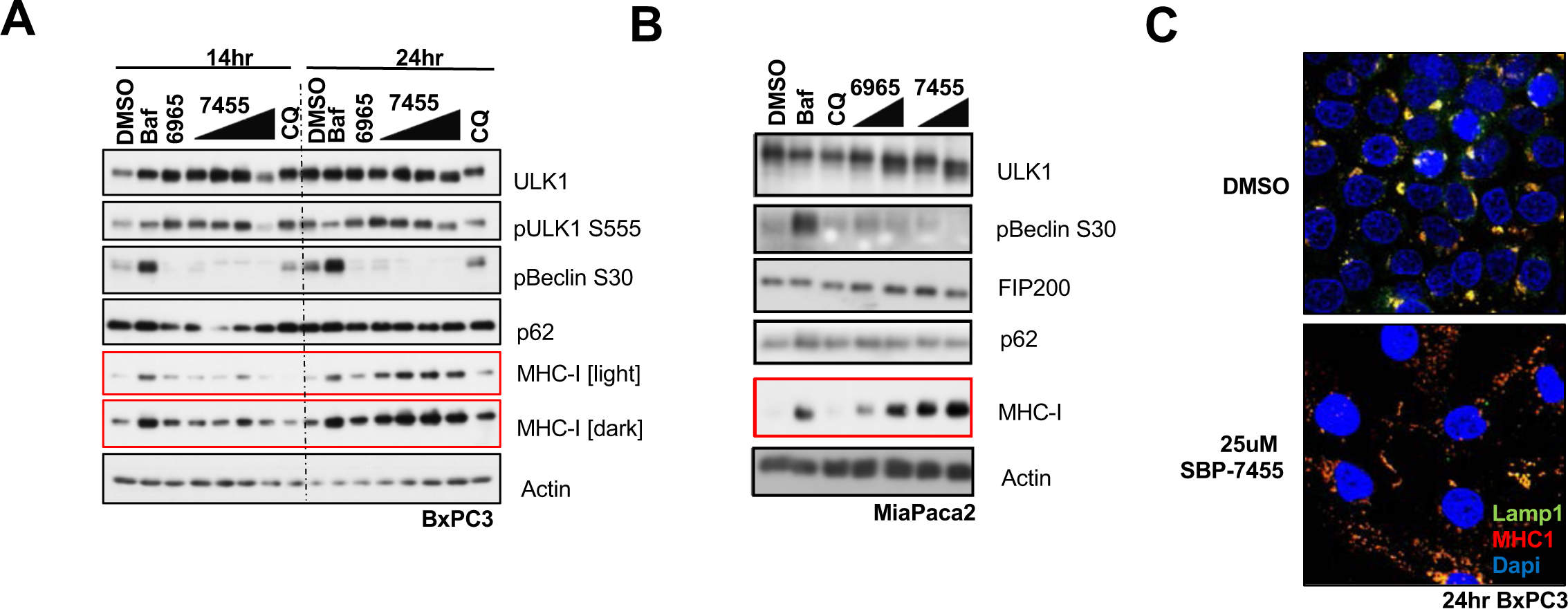
MHC-1 regulation by ULK1 inhibitors. **A**) Lysates from BxPC3 cells treated with 150nM Bafilomycin, 25μM Chloroquine, 10μM SBP-6965 and 1μM 5μM 10μM, 25μM SBP-7455 for 14 or 24 hours as indicated were analyzed by immunoblotting for MHC1 class I levels. MHC1 levels were robustly increased in response to SBP-7455 at 24hrs to a greater degree than chloroquine or bafilomycin treatment. **(B)** Lysates from MiaPaca2 cells treated with 150nM Bafilomycin, 25μM Chloroquine, 5μM and 10μM SBP-6965 and 5μM and 10μM SBP-7455 for 14 hours were analyzed by immunoblotting for MHC1 class I levels. There is clear induction of MHC1 expression in SBP-7455 samples, high dose SBP-6965, and bafilomycin treated samples. **(C)** Representative fluorescence images of MiaPaca2 cells treated with DMSO or 25μM SBP-7455 for 24hrs stained with Lamp1 (green, autophagosomes and lysosomes), MHC1 (red) and Dapi (blue, nuclei) to monitor MHC1 and Lamp1 colocalization. High degree MHC1 and Lamp1 overlap in control samples is lost after treatment with SBP-7455.

## Acknowledgements

Funding: This study was supported by grants to R.J.S. from the National Institute of Health (NIH) (R35CA220538 and P01CA120964) and to N.D.P.C. and R.J.S. from Pancreatic Cancer Action Network (PANCAN 19-65-COSF). S.N.B. was supported by NIH T32 postdoctoral training grant to the Salk Institute Cancer Center (5T32CA009370-33) and an NIH postdoctoral fellowship (F32CA206400). We thank C.M. Hung for early sgRNA design and assistance with xenograft studies.

## Author contributions

S.N.B. and R.J.S. designed the experiments and wrote the manuscript with input from all authors except as noted: S.N.B originated the project and performed experiments in Fig. 1, Fig.2A-F, Fig. 3A-D, Fig. 4, S1, S2 A-C, and S3A-F. G.S assisted with Fig. 3A-D, S2C, and S3A-F with support from M.D. and R.M.E. J.L. performed experiments in Fig 2G, Fig 3E, S2D, and S3H. A.I. assisted with Fig. 1A, 1F, Fig. 2A, S1A. A.S.L and H.R in the laboratory of N.D.P.C. provided and characterized SBP-7455 used in the study. H.T. generated and pharmacotyped the primary pancreatic organoids used in Fig 2G, Fig 3E, S2D, and S3H. D.E. supported J.L and H.T experiments and assisted with manuscript preparation.

## Data and materials availability

All data needed to evaluate the conclusions in the paper are present in the paper and/or the Supplementary Materials.

## Materials and Methods

### Cell lines and Chemicals

All cell lines were incubated at 37 °C and were maintained in an atmosphere containing 10% CO2. Miapaca2, BxPC3, and Panc1 cells were purchased from ATCC and maintained in Dulbecco’s Modified - Eagle’s Medium (MiaPaca2 and Panc1,Corning 10-013-CM) or RPMI GlutaMax (BxPC3, Invitrogen 61870-036) containing 10% fetal bovine serum (Hyclone) and 1% Penicillin-Streptomycin (Invitrogen 15140122). Cells were tested for Mycoplasma (Lonza) using manufacturer’s conditions and were deemed negative. Stable CRISPR cell lines were selected with 2μg/ml puromycin (Sigma-Aldrich P9620) and 20μg/ml Blasticidin (Invitrogen R21001) and were continuously maintained under antibiotic selection.

### Organoid culture

Detailed procedures to propagate human neoplastic pancreatic organoids have been previously described (Boj et al., 2015; Huch et al., 2013; Mihara et al., 2016; Seino et al., 2018). Organoids were propagated in Growth Factor Reduced Matrigel and Human Complete Feeding Medium: advanced DMEM/F12, HEPES 10mM, Glutamax 1X, A83-01 500nM, hEGF 50ng/mL, mNoggin 100ng/mL, hFGF10 100ng/mL, hGastrin I 0.01μM, N-acetylcysteine 1.25mM, Nicotinamide 10mM, B27 supplement 1X final, R-spondin1 conditioned media 10% final, Wnt3A conditioned media 50% final. Detailed molecular characterization of the organoids used in this study has been previously described (Tiriac et al., 2018).

### CRISPR/Cas9 Studies

Small Guide RNAs (sgRNAs) targeting human ATG7, ATG13, FIP200, and ULK1 (guide JG) were described previously (Goodwin et al., 2017). sgRNAs targeting ATG5, ULK1 (g1B), ULK2 (gA and gC) were selected using the optimized CRISPR design tool (http://crispr.mit.edu). Guides with high targeting scores and low probability of off-target effects were chosen, targeting the 1st or 2nd coding exon, or otherwise the first most common downstream exon for transcripts reported to produce multiple isoforms from searches of Uniprot or Ensembl databases. Non-targeting guide was previously described (Hollstein et al., 2019). Oligonucleotide sequences are listed in Supplemental Table S1. Oligonucleotides for sgRNAs were synthesized by IDT, annealed in vitro and subcloned into BsmbI-digested lentiCRISPRv.2-puro (Addgene 52961) or lentiCas9-Blast (ULK2 sgRNAs, Addgene 52962). Stable CRISPR knockout pools of MiaPaca2, BxPC3, or Panc1 were generated by stable integration of pLentiCRISPRv.2 and lentiCas9-Blast plasmids by lentiviral transduction with 0.45um-filtered viral supernatant supplemented with polybrene followed by selection with 2ug/ml puromycin and 20ug/ml blasticidin Validation of guide specificity and degree of knockout was assessed by Western blot of low-passage MiaPaca2, BxPC3,or Panc1 cell pools and low-passage cells KO lines were used for Cyquant, low density, and soft agar assays. Polyclonal cell pools were injected in flanks of nude mice for tumor growth studies.

### Lentiviral production

Lentiviruses made from pLentiCRISPRv.2 were produced by co-transfection of the lentiviral backbone constructs and packaging plasmids pSPAX2 (Addgene 12260) and pMD2.G (Addgene 12259). Lipofectamine 2000 (Thermo Fisher Scientific) was used as a transfection reagent at a ratio of 3:1lipofectamine/DNA. Viral supernatant was collected 48 hrs post-transfection, filtered using a o.45uM, and either immediately incubated with cells for infection or stored at four degrees for a day prior to infection.

### 2D Proliferation Assay

CRISPR polyclonal cell pools were plated into 24 well plates in triplicate at 5^E4^ cells/well and cell counts were recorded on day 5 post treatment using Cyquant assay (Life Technologies #C35011) under manufacturers conditions and emission 562nM graphed. For treatment experiments, cells were plated into 24 well plates in triplicate at 5^E4^ cells/well and the following day changed to fresh media containing either DMSO vehicle, SBP-6965 or SBP-7455. Cell counts were recorded on day 5 post treatment using Cyquant assay (Life Technologies #C35011) under manufacturers conditions.

### 2D Low Density Assay

300 MiaPaca2 or Panc1 cells were plated on 6cM dishes in full media (10% FBS in DMEM). After 10 days cells were washed with PBS, fixed for 10 mins in 10% formalin (EMD Millipore R04586-82) at room temperature, and stained for 20 mins with 0.25% crystal violet (Sigma C3886). Plates were washed, imaged using a LI-COR Odyssey CLx (Licor Inc) and Image Studio Lite software, and analysis of colony number and colony size were performed using ImageJ software.

### Soft Agar Colony Growth Assay

0.015 million MiaPaca2 cells or 0.025 million Panc1 cells were mixed with agarose in 2x growth medium (final concentration 0.3%), plated on top of a solidified layer of 0.6% noble agar in 6 well plates and fed every 4-5 days with growth medium. Cell were collected after 3 weeks, stained with 0.02% Giemsa (Sigma-Aldrich 48900) for 10mins at room temperature, then washed with PBS and stored at 4 degrees overnight for stain development. Plates were imaged using a LI-COR Odyssey CLx (LI-COR Inc) and Image Studio Lite software (LI-COR Inc.) Analysis of colony number and colony size were performed using ImageJ software (NIH).

### mRNA preparation and qPCR

For analysis of *ULK2* expression, mRNA was isolated from cells using an RNAeasy Plus Mini kit (QIAGEN Inc, Valencia, CA). One-step qRT-PCR reactions were performed in triplicate using QuantiTech RT mix (QIAGEN) on the Bio-Rad C1000 Thermocycler and CFX96 system (Bio-Rad Laboratories, Hercules, CA). Duplicate reactions were prepared without reverse transcriptase to confirm the absence of genomic DNA contamination. Relative gene expression was calculated using the ΔΔCT method and normalized to Actin. 95% confidence intervals for each sample were calculated using the sum of the squares method.

### Western blot

For biochemical analysis of tumors, tumors were immediately snap frozen in liquid nitrogen and homogenized on ice in lysis buffer. Protein lysates were equilibrated for protein levels using a BCA protein assay kit (Pierce) and resolved on 8% SDS-PAGE gels depending on the experiment. For biochemical analysis of cells, cell lysates were prepared in lysis buffer, centrifuged and equilibrated for protein levels as above. Lysates were resolved on 8-12% SDS-PAGE gels depending on the experiment.

### Pharmacotyping of organoids

Organoids were pharmacotyped following previously described methodology (Tiriac et al., 2018). In short, following single cell dissociation, 1,000 viable cells were plated per well in in 20μL 10% Matrigel / Human Complete Organoid Media. Therapeutic compounds were added 48 hours post plating using a D300E drug printer (Tecan) after the reformation of organoids was visually verified. SBP-7455 was dissolved in DMSO, Hydroxychloroquine (HCQ) was dissolved in water and a working dilution was made in DMSO. All treatment wells were normalized to 0.5% DMSO content. Single agent 10-point dose response curves were performed in triplicate for each organoid line (range from 1.0×10^−8^ M to 1.0×10^−4.3^ M for SBP-7455, 1.0×10^−7.3^ M to 1.0×10^−4^ M for HCQ). After 5 days cell viability was assessed using brightfield and phase imaging, and CellTiter-Glo as per manufacturer’s instruction (Promega) on a SparkCyto (Tecan) plate reader. A four-parameter, variable slope log-logistic function with upper limit equal to the mean of the DMSO values was used to identify the IC50 values. The synergy analyses between the MEK inhibitor Trametinib and either SBP-7455 or HCQ were performed using Combenefit (Di Veroli et al.).

### Fluorescent Autophagy Assays

#### Cyto-ID

0.08 million BxPC3 cells were plated on glass coverslips in 24well plates in duplicate in full media. The following day, wells were washed once with PBS and incubated in fresh media containing DMSO or 25μM SBP-7455 for 2hrs. After two hours, 4 μM AZD-8055 was added to the media and incubated for 24hours. Cells were then stained for autophagic vacuoles using Cyto-ID detection kit 2.0 (Enzo Life Sciences, ENZ-KIT175-0050) per the manufactures instructions and fixed for 10mins in 4% PFA. Cells were counterstained with Dapi for 5 minutes and coverslips were mounted in FluoromountG (SouthernBiothech). Images were acquired on a Zeiss Axioplan2 epifluorescence microscope coupled to the Openlab software. 5 random fields per condition were acquired using the 62Xobjective and representative images are shown.

#### Incucyte mCherry-GFP-LC3

MiaPaca2 and BxPC3 pancreatic lines were generated to stably express mCherry-GFP-LC3 construct to allow for monitoring autophagy through retroviral infection and puromycin selection as previously described (Castillo et al., 2017). 0.01 million cells were plated per well in 96 well black plates in 4-6 wells per condition. The next day, cells were treated with indicated drugs and imaged across up to 72 hours using an Incucyte S3.6-9 pictures were taken at 20x magnification per well and both well confluence and GFP/mCherry ratios were quantitated using Incucyte software (2020B). Cells that do not express mCherry or GFP were used to determine the background signal. Total integrated intensity (RCU/GCU x μm2/image) were used to calculate mCherry or GFP signal.

### Apoptosis Analysis

#### FACS

0.6 million MiaPaca2 or Panc1 cells were plated on 6cM dishes. The next day, cells were washed once with PBS and incubated in indicated concentrations of SBP-7455, 6965, chloroquine, or DMSO containing media for 72 hours. Cells were collected, washed once in PBS, trypsinized, and pelleted. Cells were then washed in 1X AnnexinV buffer and stained with AnnexinV and 7-amino-actinomycin D (7-AAD) according to the manufacturer’s protocol (BD Pharmingen, San Diego, CA). Briefly, cells were re suspended in AnnexinV buffer to a concentration of 1×10^6^/mL. Cells were then stained with 5 μL of phycoerythrin (PE)-conjugated AnnexinV antibody (Cat #556422 BD Pharmingen) and 5 μL of 7AAD (Cat# 51-2359KC BD Pharmingen) and incubated at room temperature for 15 minutes. 400 μL of AnnexinV buffer was then added to each sample with gentle mixing. Stained cells were analyzed using a FACScan flow cytometer (Becton Dickinson, San Jose, CA) and flow cytometry data was analyzed using FlowJo v.10.1 software (Tree Star Inc., Ashland, OR).

#### Western

0.6M MiaPaca2 cells were plated on 6cM dishes. The next day, cells were washed once with PBS and incubated in fresh DMEM or EBSS media containing DMSO, 500nM Bafilomycin, 20μM chloroquine, SBP-6965 (5 or 10 μM) or SBP-7455 (5 or 10 μM) for 14 hours.

### Mouse Studies

All procedures using animals were approved by the Salk Institute Institutional Animal Care and Use Committee (IACUC). For flank tumor studies, 6-8 week old female J;Nu mice were purchased from Jackson labs (007850). 5 million MiaPaca2 cells stably expressing sgRNAs targeting ULK1 (g1B) or ULK2 (g2C) or non-targeting control in 100ul of 1:1 PBS and Matrigel (BD 354234) were injected bilaterally in the flanks of the nude mice. Tumor size was measured twice a week using calipers and tumor volume calculated using the formula 0.5 ×(Length × width^2^). Once mice in the control group reached endpoint tumor volume of 2000mm^3^, mice were euthanized using CO_2_ and tumors were collected from all groups, weighed, and flash frozen in liquid nitrogen for downstream western blot analysis.

### Fluorescent Microscopy

0.08 million BxPC3 cells were plated on glass coverslips cells in 24-well tissue culture plates. The following day, wells were washed once with PBS and incubated in fresh media containing DMSO or 25μM SBP-7455 for 24hrs. Cells were the fixed in 4% PFA in PBS for 10 minutes and permeabilized in 0.2% Triton in PBS for 10 minutes. The following primary antibodies were used: mouse monoclonal MHC class 1 (Sigma H1650) and Lamp1 antibody (Cell Signaling Technologies #9091). Secondary antibodies used were anti-rabbit Alexa488 and anti-mouse Alexa594 (1:1000; Molecular Probes). Cells were then fixed and counter stained with DAPI.Coverslips were mounted in FluoromountG (SouthernBiothech). Images were acquired on a Zeiss Axioplan2 epifluorescence microscope coupled to the Openlab software. 5 random fields per condition were acquired using the 62Xobjective and representative images are shown.

### Antibodies and Reagents

For western blotting, antibodies from Cell Signaling Technologies (Denvers, MA USA) were used diluted at 1:1000 unless otherwise noted: ATG5 (#12994), ATG7 (#2631), ATG13 (#6940), ATG14 (#5504), phospho-ATG14 S29 (#92340), phospho-Beclin S30 (#54101), Beclin1 (#3495), phospho-ULK1 S757 (#6888), phosphor-ULK1 S555 (# 5869), Ulk1 (#8054), LC3B (#3868), FIP200 (#12436), ATG101 (#13492), PARP (#9542), phospho-ACC S79 (#3661), Caspase 3 (#9661), cleaved caspase 3 (#9663), phospho-TBK1 (#5483), TBK1 (#3013). Antibodies from other companies: β-actin from Sigma (A5541), phospho-FAK Y397 from Abcam (ab4803), FAK from Epitomics (2146-1), p62 SQSTM1 from Progen (03-GPP62-C), phospho-Atg13 S318 (human S355) from Rockland (600-401-C49S), NCOA4 from Santa Cruz Biotechnology (sc-373739) and MHC class 1 from Abcam (ab70328).

Reagents: EBSS (14155-063) from Gibco/Life Technologies. SBP-6965 and SBP-7455 from the Cosford Lab (Sanford Burnham Prebys). Chloroquine (C6628) and bafilomycin A1 (B1793) from Sigma. AZD-8055 (A-1008) from Active Biochem. Cisplatin (2251) and Gemcitabine (3259) from Torcis.

### Statistical analyses

Statistical analyses are described in each figure and all were performed using Graph Pad Prism 7.

---

**Supplementary Table S1.**
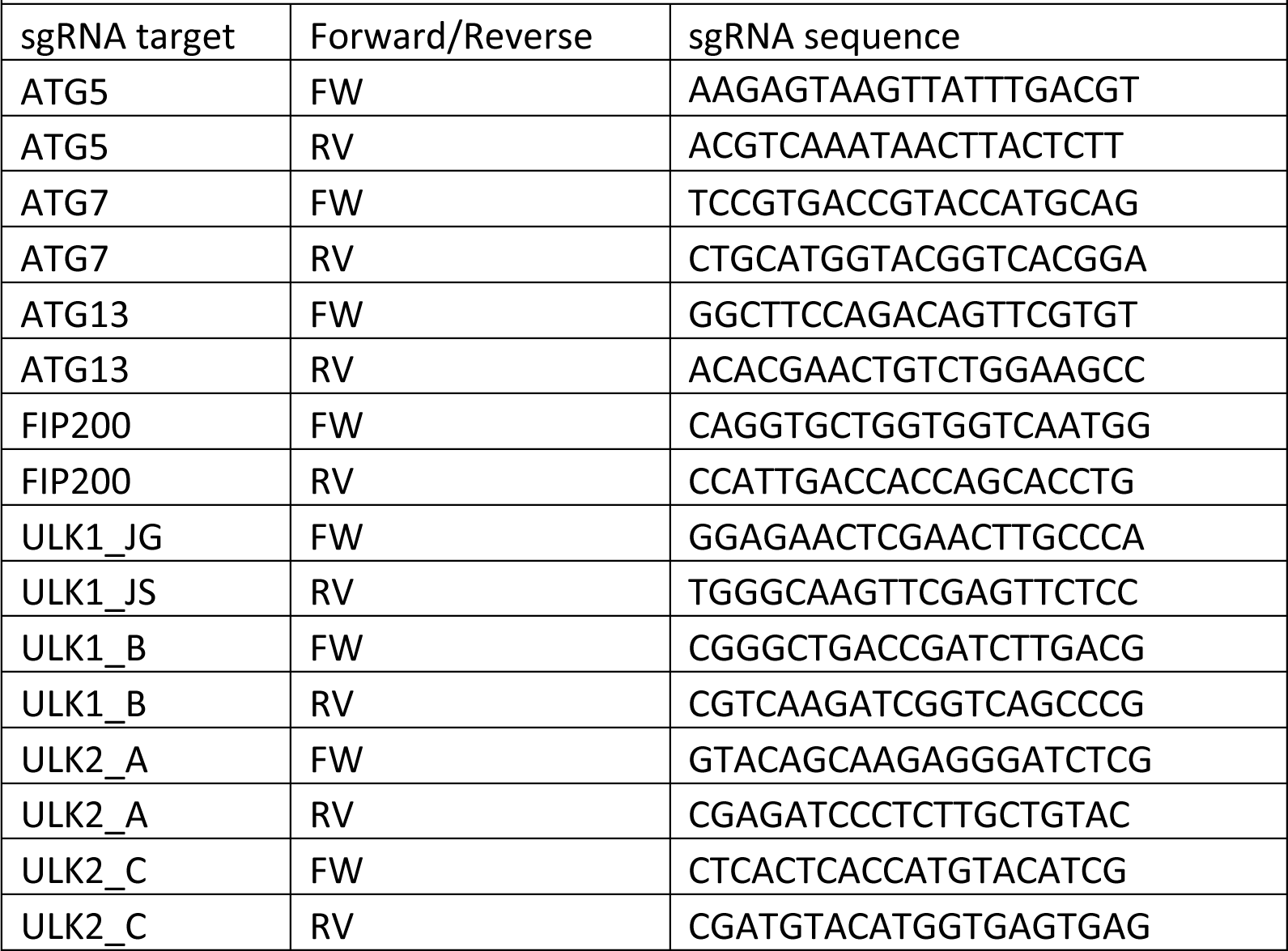

**Figure S1.**
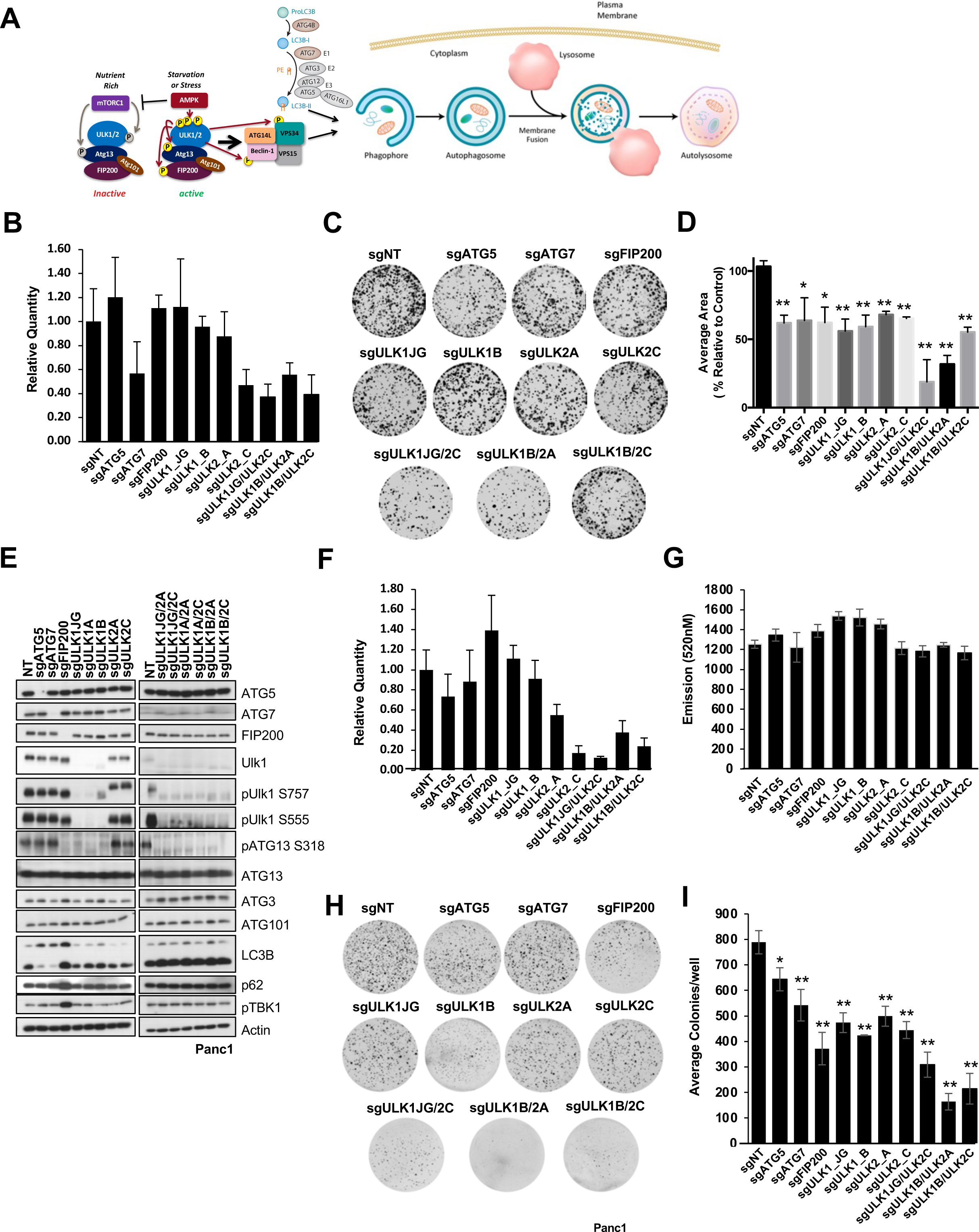
Genetic deletion of ULK1 and ULK2 impairs colony formation and soft agar growth of pancreatic tumor cells. **(A)**Schematic of the core components of the autophagy pathway with emphasis on autophagy initiation. Phosphorylation events that activate autophagy are in yellow and phosphorylation events that inhibit autophagy are in gray. **(B)** RT-qPCR analysis of Ulk2 expression in a panel of CRISPR generated MiaPaca2 cell lines reveals significant ablation of ULK2 expression in sgULK2 single and sgULK1/sgULK2 expressing lines. Data are expressed relative to control sgNT sample with 95% confidence intervals. (**C-D)** Low density growth of MiaPaca2 CRIPSR lines. (C) Representative images after cells were plated in low density conditions and grown out for 10 days. (D) Quantitation of well area covered by cells expressed as a percentage of sgNT control averaged across two independent experiments each plated in triplicate, plotted +/−SD. Low density growth is impaired by loss of all autophagy regulators including ULK1 and ULK2 and is most striking in ULK1/ULK2 double knock out cells. Significance was determined using one way ANOVA (p<0.005) and t-test (*=p<0.05, **=p>0.01) (**E)** Western blots on lysates from Panc1 cells treated with lentiviral small guide RNAs (sg) targeting autophagy regulators ATG5, ATG7, FIP200, ULK1 or ULK2 alone, or lines targeted to co-delete ULK1 and ULK2 demonstrate efficient deletion of protein targets. (**F**) RT-qPCR analysis of Ulk2 expression in a panel of CRISPR generated Panc1 cell lines reveals significant ablation of ULK2 expression in sgULK2 single and sgULK1/sgULK2 expressing lines. Data are expressed relative to control sgNT sample and confidence intervals were generated using the sum of the route of the squares method.**(G)** CRISPR panel of Panc1 cells were plated in adherent growth conditions and cell number after 7 days was quantified using cyquant in two independent experiments plated in triplicate. Loss of autophagy regulators including ULK1 and ULK2 do not significantly alter 2D growth of Panc1 calls. (**H-I)** Soft Agar analysis of Panc1 cell lines. (H) Representative images after 3 weeks of growth. Soft agar growth is most impaired after loss of both ULK1 and ULK2. (I) Quantitation of average colony number per well expressed as a percentage of sgNT control cell colony number plated in triplicate +/−SD. Data representative of two independent experiments. Significance was determined by t-test (*=p<0.05, **=p>0.01)

**Figure S2.**
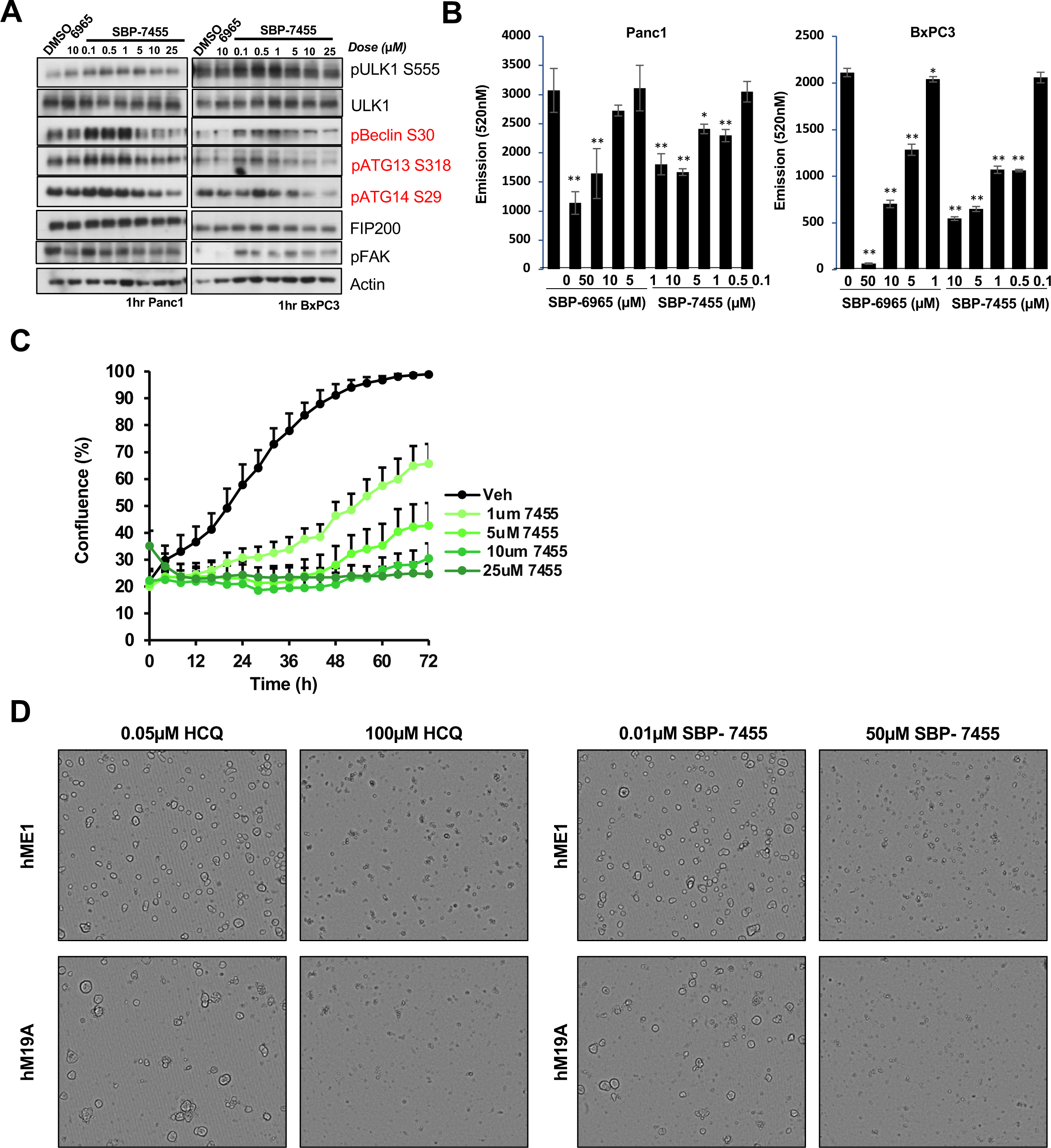
ULK1/ULK2 inhibitors impair growth of multiple pancreatic tumor lines and organoids in vitro. **A**) Lysates from Panc1 and BxPC3 cells treated with ULK1 inhibitors SBP-6965 and SBP-7455 at indicated concentrations for 1 hour were analyzed by immunoblotting for ULK1 inhibition and off-target signaling. SBP-7455 effectively inhibited phosphorylation of Ulk1 substrates Beclin, ATG13, and ATG14. **(B)** Dose dependent inhibition of MiaPaca cell growth by Ulk1 inhibitors SBP-6965 and SBP-7455 after 72hrs of treatment demonstrated using Cyquant assay run in triplicate in three independent experiments (representative run plotted +/− SD). SBP-7455 inhibits cell growth at lower concentrations than SBP-6965. Significance relative to DMSO control was determined using t-test (*=p<0.05, **=p>0.01). **(C)** BxPC3 cells stably expressing LC3-mCherry-GFP construct were treated with DMSO or SBP-7455 at indicated concentrations and well confluence monitored over 72hrs by Incucyte brightfield imaging and quantification of six wells per condition +/− SD, showing dose dependent inhibition of cell growth. **(D)** Representative images of hME1 and hM19A organoids treated with hydroxychloroquine (HCQ, 0.05μM/ 100μM) or SBP-7455 (0.01μM/ 50μM) as indicated for 5 days. High doses of both HCQ and SBP-7455 significantly impair organoid growth and survival.

**Figure S3.**
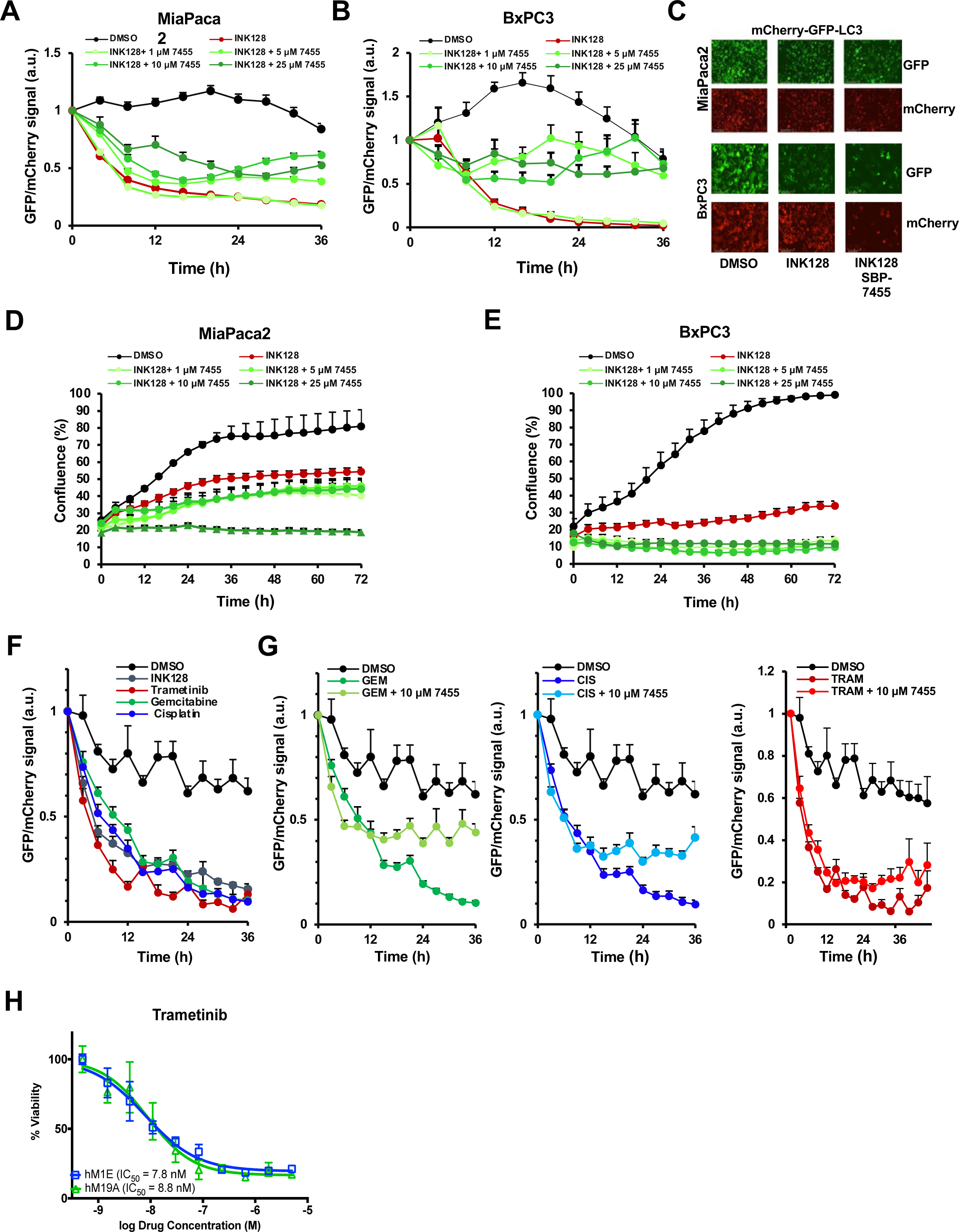
Time-lapse analysis of autophagic flux and cell growth. **(A-B)** Full 36hr timecourses of GFP/mCherry ratios in MiaPaca2 (A) and BxPC3 (B) cells stably expressing mCherry-GFP-LC3 construct treated with DMSO or SBP-7455 for 2 hours then 1μM INK128 (mTOR inhibitor) added in combination with ULK1 inhibitor to monitor autophagy flux +/− SEM, corresponding for Figure A,B. **(C)** Representative images of DMSO, 1 μM INK128, and combination of 1 μM INK128 with 10μM SBP-7455 treated Miapaca2 (top) and BxPC3 (bottom) cells expressing mCherry-GFP-LC3 at 24 hours taken at 20x magnification. **(D-E)** MiaPaca2 (C) and BxPC3 cells stably expression mCherry-GFP-LC3 construct treated as in (A-B) were monitored for well confluence monitored over 72hrs by Incucyte brightfield imaging and quantification of six wells per condition (22-36 pictures per condition +/− SEM). **(F)** Full 36 hour time courses of GFP/mCherry ratios in mCherry-GFP-LC3 expressing MiaPaca2 cells treated with indicated drugs +/− SEM corresponding to Figure 3D **(G)** Full 36 hour timecourses of GFP/mCherry ratios in mCherry-GFP-LC3 expressing MiaPaca2 cells treated with indicated drugs +/− SEM corresponding to Figure 3E. **(H)** Two human organoid lines (hM1E and hM19A) were treated with 10 concentrations of Trametinib for 5 days and dose response curves generated. Both lines responded in a dose-dependent manner to Trametinib with an IC50 of 7.8n M and 8.8nM, respectively.

